# Loss of Calcitonin Gene Related Receptor component protein (RCP) in nervous system can bias “gepant” antagonism

**DOI:** 10.1101/2024.10.25.620369

**Authors:** Shafaqat M. Rahman, Ian Dickerson, Anne E. Luebke

**Affiliations:** University of Rochester, Department of Biomedical Engineering, Rochester, NY 14627; University of Rochester Medical Center, Department of Neuroscience, Del Monte Institute of Neuroscience, Rochester, NY 14642

**Author notes:** Corresponding author: Anne E. Luebke.

## Abstract

We examined calcitonin gene-related peptide (CGRP)’s effects on behavioral surrogates for motion-induced nausea and static imbalance in the nestinRCP (-/-), a novel mouse model that loses expression of receptor component protein (RCP) in the nervous system after tamoxifen induction. The assays used were the motion-induced thermoregulation and center of pressure (CoP) assays. Findings suggest CGRP’s affects behavioral measures in the nestinRCP (-/-) similarly to littermate controls, since CGRP was observed to increase female sway and diminishes tail vasodilations to provocative motion in both sexes. However, the CGRP-receptor antagonist olcegepant did not antagonize CGRP’s effects in the nestinRCP (-/-), whereas it was effective in littermate controls. Findings suggest RCP loss may change the sensitivity of the CGRP receptor and affect the efficacy of receptor antagonists.

**Significance Statement:** Research in calcitonin gene-related peptide (CGRP) has primarily focused on ligand-receptor interactions at the calcitonin-like receptor (CLR) and receptor activity-modifying unit 1 (RAMP1) subunits. However, the role of receptor component protein (RCP), which mediates signaling via the Gα-stimulatory pathway, is less understood. A novel tamoxifen-inducible mouse model, nestinRCP (-/-), was generated to study loss of RCP in CGRP signaling in the nervous system, and behavioral changes to motion-induced nausea and postural sway were studied after systemic injections of CGRP or CGRP co-delivered with migraine drugs. Findings from this study suggest the loss of CGRP-RCP can bias “gepant” antagonists like olcegepant, and may promote development of therapies to inhibit the RCP-CLR interactions.

## Introduction

Vestibular migraine is a migraine type where patients have a history of migraine complications - like sensory hypersensitivities and headache pain - but also develop a maladaptive sensitivity to motion that leads to recurrent vertigo and feelings of dizziness[1]. In challenging environments, VM patients may exhibit postural sway and a susceptibility to motion sickness and nausea. Much like migraine, VM is characterized by a female preponderance [2–5].

Calcitonin gene-related peptide (CGRP) is a neuropeptide and neuromodulator with an emphasized role in migraine pathophysiology [6–8]. In the last several decades, abundant translational research has led to the development of therapies that inhibit CGRP signaling for migraine treatment. These therapies include the small molecule receptor antagonists – the ‘gepants’ – as well as monoclonal antibodies that block either the CGRP peptide or the receptor [9]. It is expected that by the year 2027, millions of people with migraine will be using CGRP blockers in the treatment of their migraine [10].

The calcitonin gene-related peptide (CGRP) receptor is composed of three subunits: a G-protein coupled receptor (GPCR) called calcitonin receptor-like receptor (CLR), a transmembrane subunit called receptor activity-modifying protein 1 (RAMP1), and a cytosolic, peripheral membrane subunit called receptor component protein (RCP) [11]. Abundant research has been done to characterize CLR/RAMP1 interactions in CGRP signaling, but much work is still required to elucidate the role of RCP.

RCP is 148 amino acids and 17 kDa in size. While RCP is not needed for binding of the CGRP peptide to the CGRP receptor, and is not needed for trafficking of CLR and RAMP1 to the membrane surface, RCP is a key aspect of CGRP signaling as it stabilizes CLR interaction with the Gα stimulatory protein (Gαs) and mediates downstream signaling that leads to intracellular cAMP accumulation [12]. RCP protein expression correlates with CGRP efficacy and the signaling of related peptides – like adrenomedullin, which also relies on RCP [11]. RCP is believed to act as a positive allosteric modulator and promote agonist signaling bias at the CGRP receptor and related B family GPCRs [13].

CGRP and its receptor components are widely present in the nervous system. RCP is expressed broadly in the brain, spinal cord, smooth muscle and cerebral vasculature.

Immunostaining has shown RCP to be in the nucleus of motor neurons and cerebellar Purkinje cells of rats, and the activation of the CGRP receptor is hypothesized to exert a role in cerebellar plasticity [14].

There is also abundant evidence suggesting CGRP and its receptor are involved in inner ear function and vestibular sensation. CGRP is widely distributed throughout the vestibular system, is involved in the vestibulo-ocular reflex, has a role in otolith timing dynamics, and is found on vestibular efferent fibers that innervate the inner ear [15–17]. Formation of functional CGRP receptors is associated with sound-evoked nerve activity, as cochlear function develops with the increased formation of CLR/RAMP1/RCP receptors in the mouse cochlea [18]. In addition, we have shown that systemically delivered CGRP at 0.1 mg/kg can modulate the auditory brainstem response and vestibular sensory-evoked potentials of mice, suggesting CGRP can act directly on the inner ear and provides a basis for its involvement in neurological disorders such as vestibular migraine. We have also shown that systemic CGRP also influences mouse surrogate behaviors for imbalance, phonophobia, and motion-induced nausea [19, 20].

Up until now, it is understood that CGRP receptor expression is ubiquitous in the nervous system, and that RCP is a crucial to cAMP mediated CGRP signaling. However, it is unclear what effects would arise when RCP is depleted *in vivo* in neurons and glia, and how this change in signal processing impacts physiological and behavioral outcomes. We hypothesized that neural loss of RCP will modify CGRP signaling and result in changes in surrogate behaviors that rely on vestibular cues. We also hypothesized that migraine drugs that attenuate CGRP signaling may change in their efficacy with the selective loss of RCP.

In order to determine the effects of RCP loss in the nervous system and study changes in mouse behavior, we created a novel mouse model called the nestinRCP (-/-) that relies on the tamoxifen-inducible creER system. These mice age and develop similarly as wildtype, but the introduction of tamoxifen leads to directed knockout of RCP expression in cells governed by the nestin promoter, which are primarily cells of the nervous system.

NestinRCP (-/-) and tamoxifen-treated control mice were studied for changes in motion-induced nausea and postural sway using the motion-induced thermoregulation and center of pressure (CoP) assays previously studied in wildtype C57BL6/J, found in [19, 20]. The CGRP- receptor antagonist olcegepant and the selective serotonin receptor (5HT1B/1D) agonist rizatriptan were used to study the efficacy of migraine blockers in the nestinRCP (-/-) and controls.

## Methods

### Animals

We have generated mice with a loxP conditional knockout of the 2nd exon of the *Crcp* gene. We then crossed these *Crcp*-loxP mice with nestin-Cre^ER^ mice, resulting in mice with blocked neuronal RCP expression following tamoxifen induction (*nestinRCP(-/-)*). In parallel, control mice included nestin-cre^ER^ mice that did not express *Crcp*-loxP and *Crcp*-loxP mice not expressing Cre. The nestin-Cre^ER^ mice without *Crcp*-loxP were tested to ensure Cre production did not produce confounding behavioral effects. No differences were detected in the behaviors between controls after tamoxifen treatment, so they were grouped together as tamoxifen-treated controls. Mice were placed under the care of the University Committee on Animal Resources (UCAR) at the University of Rochester. Mice were housed with ad libitum access to necessities (food, water, bedding, and enrichment). A total of 64 nestinRCP (-/-) mice (31M/31F) and 67 control mice treated with tamoxifen (34M/33F) were tested. Mice were equilibrated before testing at an ambient temperature between 22-23°C for at least 30 minutes, and mice were aged around 3 to 7 months. Different cohorts of mice were used to test motion-induced nausea and postural sway. Testing occurred from 9:00 am to 5:30 pm. Mice screened for instances of alopecia were excluded from the motion-induced thermoregulation experiments.

### Tamoxifen Induction

Intraperitoneal injections of tamoxifen were performed using Jackson Laboratory’s protocol, previously validated in ubiquitous Cre/ER models. In summary, tamoxifen (Sigma-Aldrich) was prepared in corn oil at a concentration of 20 mg/ml by overnight shaking at 37°C. Mice were injected daily at 75 mg/kg tamoxifen (in 100 uL doses) once every 24 hours for four days. Mice began behavioral experiments approximately 10 days after the fourth injection.

### Drug administration

All intraperitoneal (IP) injections were performed with a fine 33-gauge insulin syringe.

Dulbecco PBS (saline) served as vehicle control diluent. The CGRP-receptor antagonist olcegepant and the selective serotonin receptor agonist rizatriptan were used as migraine blockers in this study. The concentrations are listed: 0.1x, 0.5x, and 1.0x CGRP were prepared at 0.01, 0.05, and 0.1 mg/kg (rat ɑ-CGRP, Sigma), 1.0x olcegepant was prepared at 1.0 mg/kg (BIBN4096, Tocris), and 1.0x rizatriptan was prepared at 0.6 mg/kg (Sigma-Aldrich). Mice were tested approximately twenty to thirty minutes after injections. Animals were gently handled without the need for anesthesia. All animal procedures were approved by the University of Rochester’s (IACUC) and performed in accordance with the standards set by the NIH.

### Center of Pressure (CoP) testing for postural sway

Postural sway was examined using the center of pressure (CoP) test which acts as a mouse surrogate behavior for static imbalance, using the protocol we used here [19].

Approximately 10-12 CoP areas were captured per mouse, and a 10% robust outlier removal (ROUT) was used on an individual mouse’s trials to remove outlier CoPs. After this exclusion, each mouse - at minimum - had six different CoP areas to average for a given testing condition. Individual mice were not excluded in this study (**Fig. 1B**). For sway, mice were repeatedly tested every 2 to 4 days when examining effects of CGRP and migraine blockers.

**Fig. 1:**
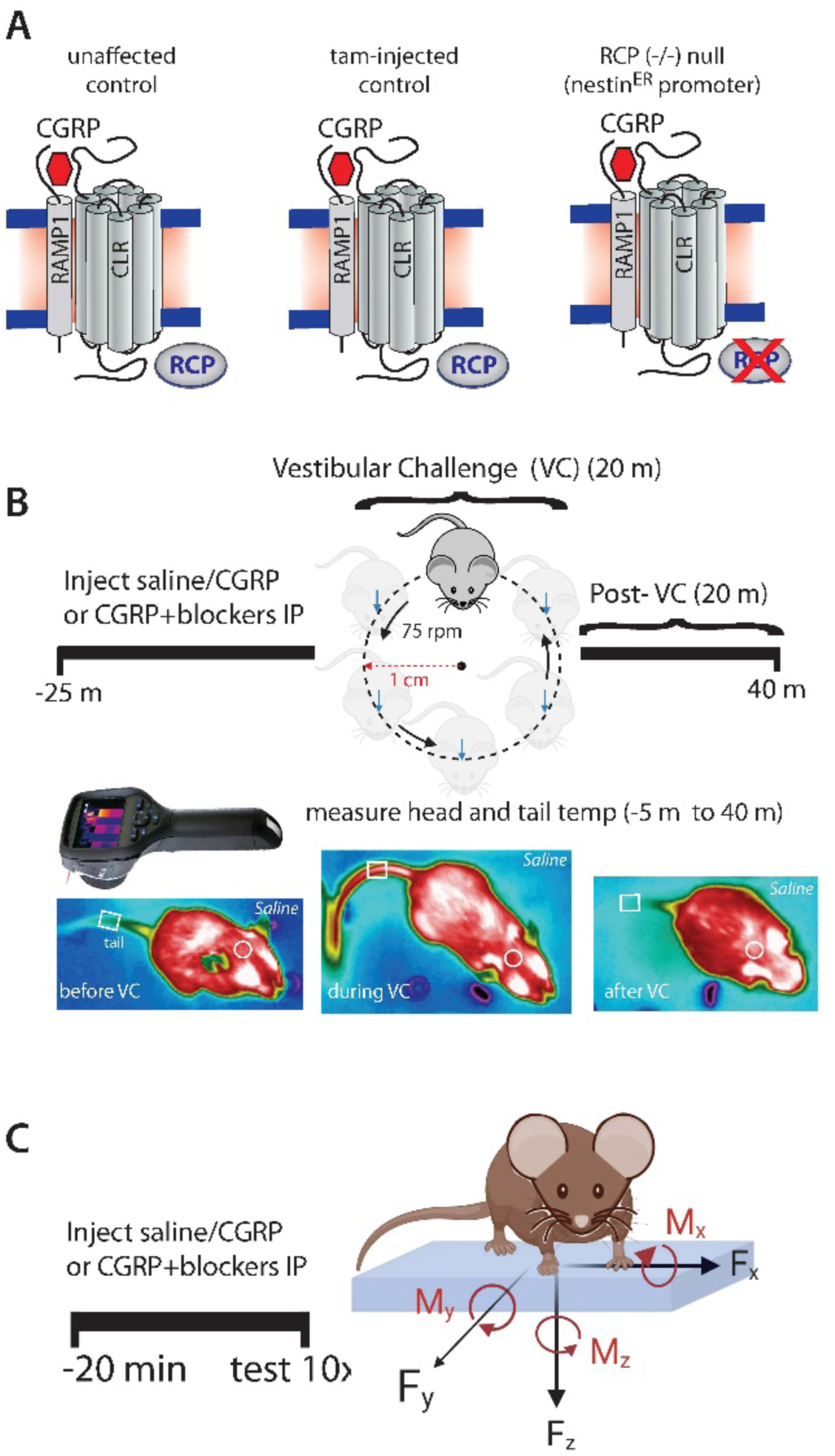
The CGRP receptor consists of calcitonin-like receptor (CLR), receptor activity modifying protein 1(RAMP1), and receptor component protein (RCP). (A) Tamoxifen-treated control mice do not have changes in their receptor structure from unaffected controls. The nestinRCP (-/-) mice lose receptor component protein (RCP) due to tamoxifen induction. **(B-C)** Methods for motion-induced thermoregulation as a surrogate behavior for motion-induced nausea and center of pressure (CoP) testing as a surrogate behavior for postural sway are shown.

### Motion-Induced Thermoregulation for assessing motion-induced nausea

We used a FLIR E60 IR camera (model: E64501) to measure the temperatures of the head and tail of C57B6/J mice. The measurements were taken for a total of 45 minutes, following a protocol we previously used [20]. In summary, we recorded baseline measurements for five minutes before initiating the provocative motion (-5 ≤ t ≤ 0). Mice were then recorded for 20 minutes (0 ≤ t ≤ 20) during an orbital rotation (75 rpm, 2 cm displacement). After the rotation, mice were recorded for an additional 20 minutes to assess the recovery of temperatures to baseline (20 ≤ t ≤ 40). Data was analyzed using FLIR Tools+.

We defined Δ tail vasodilations (°C) as the transient increases in mouse tail temperature in response to the motion. These were computed by subtracting the baseline tail temperature at time t = 0 minutes from the maximum tail temperature measured during the first 10 minutes of the rotation (0 ≤ t ≤ 10). Additionally, we calculated the magnitude of hypothermia (Δ heads) by subtracting the baseline head temperature at time t = 0 minutes from the minimum head temperature recorded during the entire experiment (0 ≤ t ≤ 40).

In this assay, mice were repeatedly tested every 4 to 6 days when examining CGRP and migraine blocker effects.

### Data analysis and statistics

All statistical analyses were conducted in GraphPad Prism 9.5. Analyses were conducted separately in females and males. Multivariate ANOVAs and mixed-effect models were used to analyze Δ tail vasodilations and center of pressure ellipse areas, and are further elaborated in the results. Group averages of Δ tail vasodilations and CoP ellipse areas are reported as mean ± SEM, and significance was set at p < 0.05 for all analyses.

Analysis of Δ tail vasodilations involved setting a threshold of 1.5°C upon the data to make it a binary outcome measure. Tail temperature changes equal to or greater than +1.5°C were designated a Δ tail vasodilation and those less than +1.5°C did not meet the criteria and were labeled as diminished tail vasodilations. Mice were excluded from further testing if their Δ tail vasodilation after vehicle (saline) testing did not meet the criteria.

Second-order curve fitting was used for fitting head temperatures (B2*X^2^ + B1*X + B0) and R^2^ fit were computed per curve. Head recovery (mins) was approximated by normalizing head temperatures so that baseline head temperature at x = 0 mins started at y = 0 °C. An x-intercept quadratic model was used to approximate the recovery time, with a constraint imposed upon detecting x-intercepts passed 20 minutes (when rotation stopped).

## Results

### Tamoxifen induction does not impact the presentation of tail vasodilations or modify baseline sway in mice

Prior to assessing CGRP’s effects, we examined motion-induced thermoregulation and center of pressure (CoP) results in mice before and after they were treated with the tamoxifen induction protocol. Mice were examined for pre-tamoxifen (pre-tam) behaviors at most 10 days before the first tamoxifen injection, whereas experiments post-tamoxifen (post-tam) occurred 10 days after the fourth tamoxifen injection. No differences were observed in Δ tail vasodilations pre and post tamoxifen during the motion-induced nausea test for either tamoxifen-treated controls and the nestinRCP (-/-) **Supplementary Fig. S1A-B**). In addition, a subset of nestinRCP (-/-) mice were examined for postural sway after vehicle administration pre and post tamoxifen induction, and no differences were observed (**Supplementary Fig. S1C**).

### No difference in hypothermia magnitude between nestinRCP (-/-) and tamoxifen-treated controls

Head temperatures are depicted after intraperitoneal (IP) delivery of vehicle (saline), 1.0x CGRP, 1.0x CGRP + 1.0x Olcegepant, and 1.0x CGRP + 1.0x rizatriptan in males and females (**Fig. 2A-H**). The magnitude of hypothermia was computed during an animal’s test – designated as the Δ head – but no differences in hypothermia magnitude were observed in mice regardless of treatment (**Fig. 2I-J**).

**Fig. 2:**
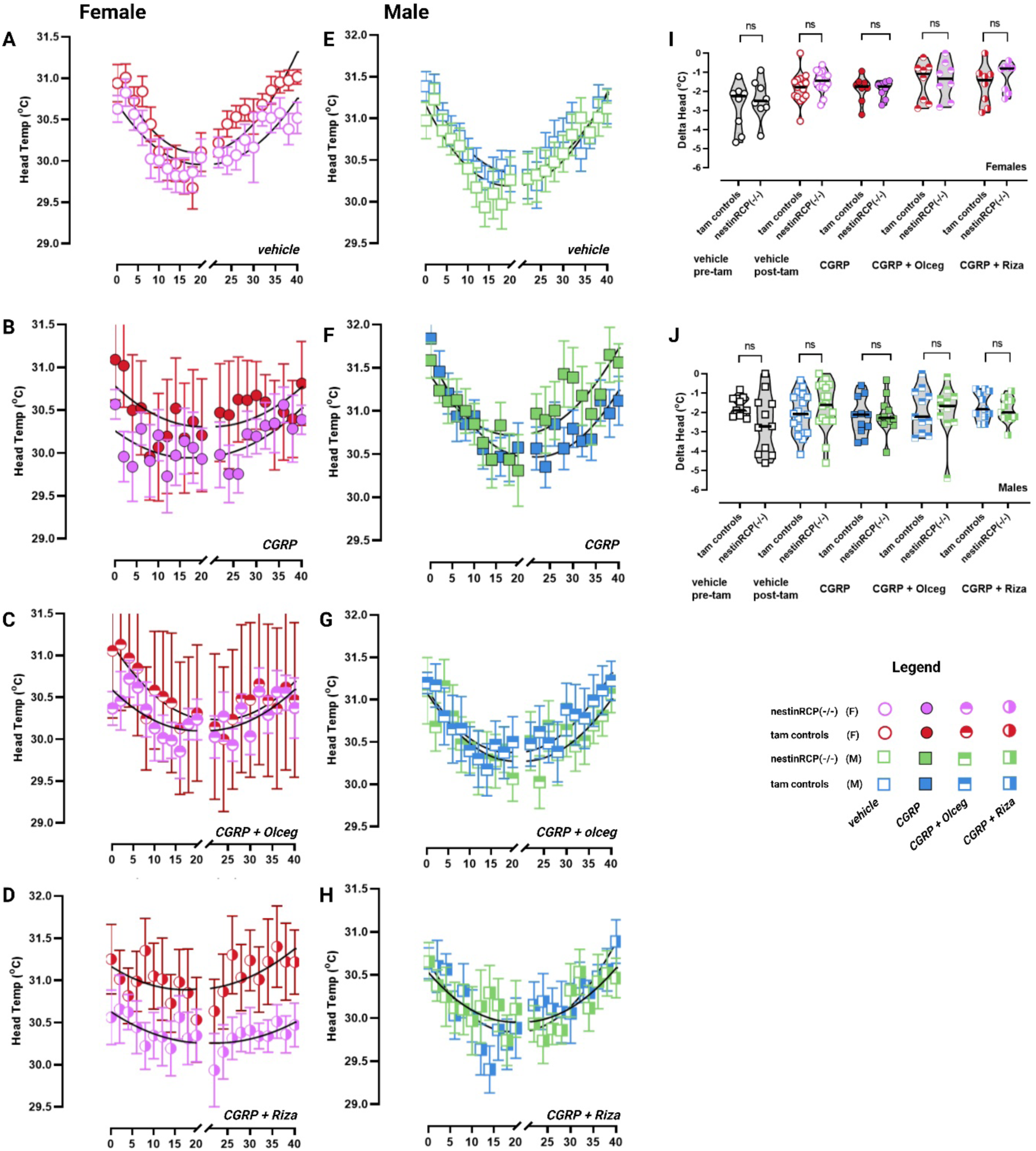
Head temperatures are shown after treatment of vehicle, CGRP, CGRP + olcegepant, or CGRP + rizatriptan in females **(A – D)** and males **(E-H).** Legend indicates symbol and color code across strain, sex, and treatment. **(I-J)** Magnitudes of hypothermia (Δ heads) were computed but no differences were seen between tamoxifen-treated controls and nestinRCP (-/-) in either sex. Sample sizes are listed: vehicle – 18M/15F nestinRCP (-/-) and 18M/15F controls; CGRP, CGRP + olceg, CGRP + riza – 11M/8F nestinRCP (-/-) and 11M/8F controls.

### Hypothermia recovery changes in male controls may reflect blocker efficacy, but neural RCP loss effects are not clear via hypothermia

At baseline, male nestinRCP (-/-) mice recover from hypothermia approximately 4 minutes faster than male controls (*F (DFn, DFd) = 10.91 (1, 36), p = 0.002*). After IP CGRP delivery, this discrepancy between male nestinRCP (-/-) and controls changed to a 7-minute difference in recovery times, with male controls taking much longer to recover from the effects of CGRP and provocative motion (*F (DFn, DFd) = 10.91 (1, 36), p = 0.002).* The treatment of CGRP+olcegepant and CGRP + rizatriptan countered CGRP – induced changes in male controls, by reducing their recovery times.

Female mice tested after vehicle and after 1.0x CGRP followed similar patterns as male mice. Female nestinRCP (-/-) mice recover from hypothermia 2.0 minutes earlier than female controls, and this discrepancy becomes a 5.0-minute difference after 1.0x CGRP, with female controls having longer recovery. However due to poor curve fits in female mice, the analysis was deemed not significant, which may be attributed to insufficient power.

The recovery times between male nestin RCP (-/-) and controls after CGRP + blockers did not differ. However, CGRP + blocker effects in the nestinRCP (-/-) complicate interpretations. CGRP + olcegepant and CGRP + rizatriptan increased recovery times in male nestin RCP (-/-) by approximately 3 minutes compared to their vehicle response, and an interpretation for this result is difficult to formulate. Nonetheless, a detailed breakdown of recovery times and curve fits is found in **Table 1**.

**Table 1.** Time to recover from hypothermia (minutes) was analyzed between tamoxifen-treated controls and nestinRCP (-/-) mice using an x-intercept quadratic model. Males and females were separated. A constraint to the model was applied so that the x-intercept analyzed occurred after time = 20 minutes. While F-statistics and p-values are listed, comparisons are made more confidently in the male curves since fit values (R^2^) are ≥ 0.60, whereas female fit curves after CGRP or CGRP + blockers poorly fit the data. Sample sizes (n’s) are listed that went into each curve fit: vehicle – 18M/15F nestinRCP (-/-) and 18M/15F controls; CGRP, CGRP + olceg, CGRP + riza – 11M/8F nestinRCP (-/-) and 11M/8F controls.

| Recovery from Hypothermia - tam controls vs nestinRCP (-/-) |  |  |  |  |  |  |  |  |
| --- | --- | --- | --- | --- | --- | --- | --- | --- |
| Sex | Treatment | tam controls |  | nestinRCP (-/-) |  | Recovery Difference | F(DFn/ DFd) | p-val |
|  |  | Recovery | Fit | Recovery | Fit |  |  |  |
| Male | vehicle | 41.00 | 0.82 | 36.60 | 0.83 | 4.4 | 10.90 (1, 36) | 0.002 |
|  | 1.0x CGRP | 46.00 | 0.85 | 38.30 | 0.63 | 7.7 | 10.90 (1, 36) | 0.002 |
|  | 1.0x CGRP + 1.0x Olceg | 38.60 | 0.83 | 41.30 | 0.68 | -2.7 | 2.45 (1, 36) | 0.13 |
|  | 1.0x CGRP + 1.0x Riza | 37.29 | 0.73 | 41.14 | 0.60 | -3.9 | 3.75 (1, 36) | 0.07 |
| Female | vehicle | 36.30 | 0.70 | 38.30 | 0.73 | -2.0 | 1.50 (1,36) | 0.24 |
|  | 1.0x CGRP | 46.00 | 0.31 | 40.90 | 0.44 | 5.1 | 1.15 (1, 36) | 0.29 |
|  | 1.0x CGRP + 1.0x Olceg | 34.72 | 0.44 | 30.19 | 0.38 | 4.5 | 2.20 (1,36) | 0.15 |
|  | 1.0x CGRP + 1.0x Riza | 36.89 | 0.35 | 42.02 | 0.43 | -5.1 | 1.64 (1,36) | 0.21 |

### Strong majority of mice exhibit disrupted nausea response at highest CGRP dose (0.1 mg/kg)

Δ tail vasodilations were examined in the nestinRCP (-/-) and tamoxifen-treated controls after administration of 0.1x, 0.5x, and 1.0x CGRP. Raw tail temperatures are depicted during the experiment after intraperitoneal injections of vehicle (**Fig. 3A, 3D**) and 1.0x CGRP (**Fig. 3B, 3E).** 2-way mixed effects analyses assessed the factors i) CGRP dose and ii) controls vs neural RCP loss. Analysis was separately done in males and females. Dunnett’s multiple comparisons test were used to compare specific dose responses to vehicle.

**Fig. 3:**
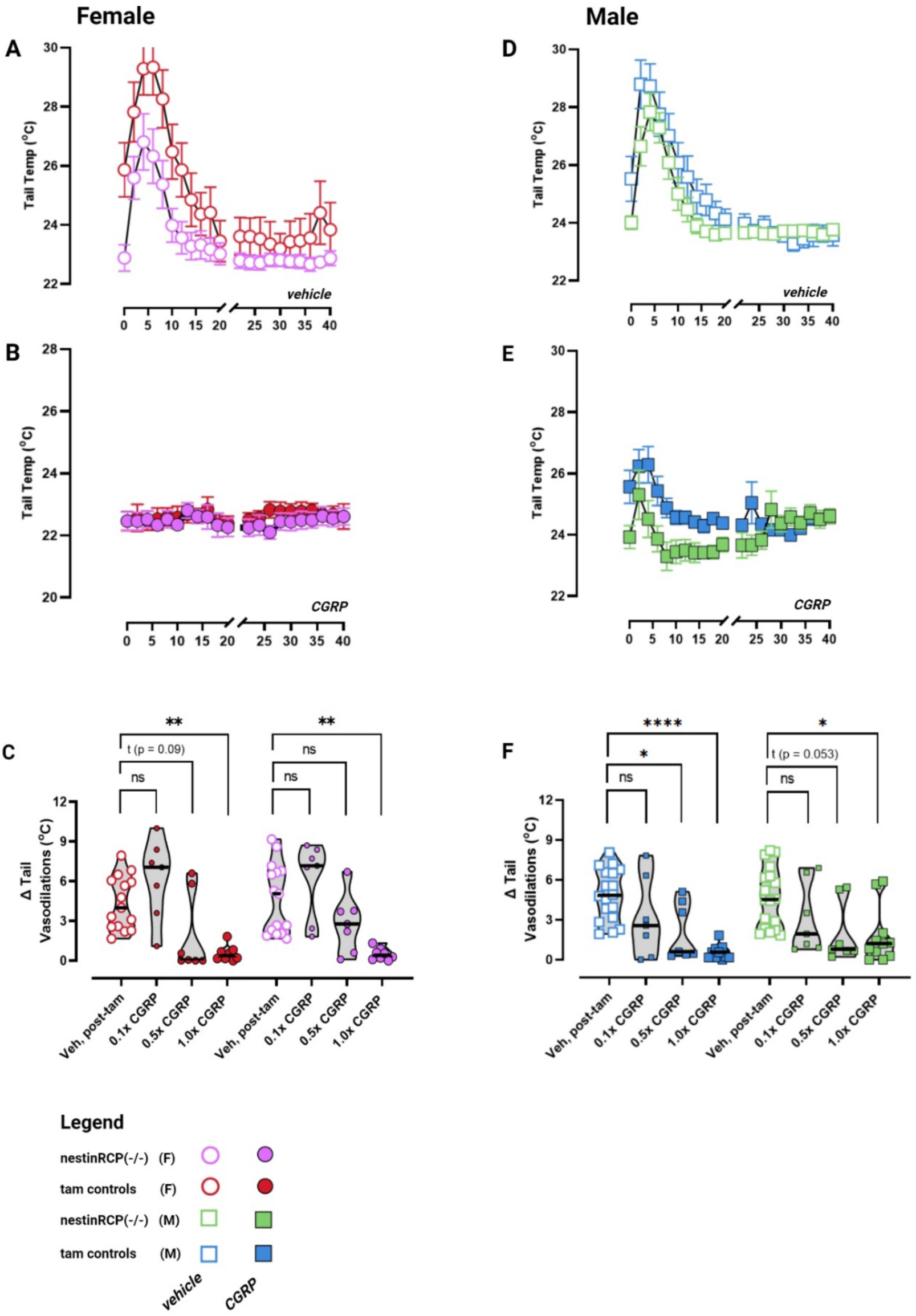
Tail temperatures and dose response analysis of Δ tail vasodilations are depicted in females (left) and males (right). **(A, D)** Mice exhibit a robust and transient tail vasodilation after the rotation starts, but this response is diminished **(B, E)** after 1.0x CGRP. **(C)** A significant difference is seen in female tamoxifen-treated controls (*adj. p = 0.004*) and female nestinRCP (- /-) (*adj. p = 0.005*) after 1.0x CGRP. **(F)** Likewise, a significant difference is seen in male tamoxifen-treated controls (*adj. p < 0.0001*) and male nestinRCP (-/-) (*adj. p = 0.048*) after 1.0x CGRP. Sample sizes are listed: vehicle – 18M/15F nestinRCP (-/-) and 18M/15F controls; 0.1x and 0.5x CGRP -7M/7F nestinRCP (-/-) and 7M/7F controls; 1.0x CGRP - 11M/8F nestinRCP (-/-) and 11M/8F controls. Full list of F-statistics and p-values can be found in middle section of Table 2. p < 0.05*, p < 0.01**, p < 0.001***, p < 0.0001****.

**Table 2.**
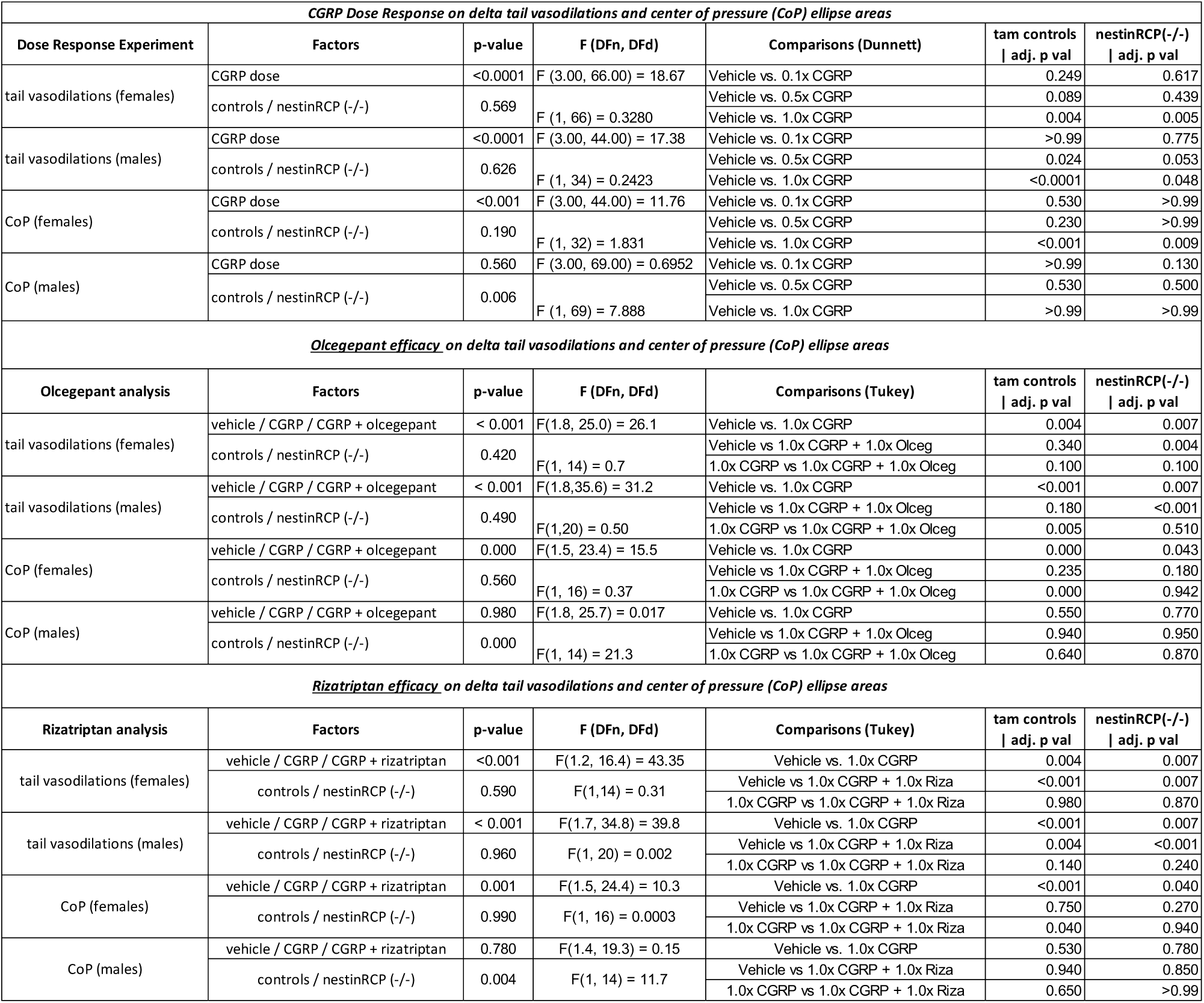
Statistics used in study are listed with F and p values. Dose responses were analyzed with 2-way repeated measures (RM) mixed effects models to handle missing values since not all animals were treated with every CGRP dose, whereas blocker studies used 2-way RM ANOVAs.

CGRP dose impacted the presentation of Δ tail vasodilations in female mice (*F (3.000, 66.00) = 18.67, p < 0.0001*), but neural RCP loss had no effect (F (1, 66) = 0.3280, p = 0.57). This CGRP dose effect was also seen in males (*F (3.000, 44.00) = 17.38), p < 0.0001*) but not neural RCP loss (*F (1, 34) = 0.2423, p = 0.63*).

Intraperitoneal delivery of 0.1x CGRP did not impact presentation of Δ tail vasodilations.

While more mice lose a robust presentation of tail vasodilations to the orbital rotation at 0.5 CGRP, a significant decrease in the presentation of Δ tail vasodilations was observed after 1.0x CGRP delivery in both sexes of nestinRCP (-/-) and tamoxifen-treated controls (**Fig. 3C, 3F**).

### Olcegepant blocks CGRP-induced changes on Δ tail vasodilations in controls, but not in the nestinRCP (-/-)

Mice were examined for the presence of Δ tail vasodilations after co-administration of 1.0x CGRP with 1.0x olcegepant (**Fig. 4A-B**). In females, a majority of tamoxifen-treated controls regained their Δ tail vasodilation when treated with olcegepant (**Fig. 4C**). However, female nestinRCP (-/-) mice did not respond to olcegepant and still exhibited diminished Δ tail vasodilations that resembled their 1.0x CGRP testing. A similar observation was seen in males (**Fig. 4D**). A majority of male controls regained their Δ tail vasodilation due to olcegepant, but the nestinRCP (-/-) males did not recover with this CGRP-receptor antagonist.

**Fig. 4:**
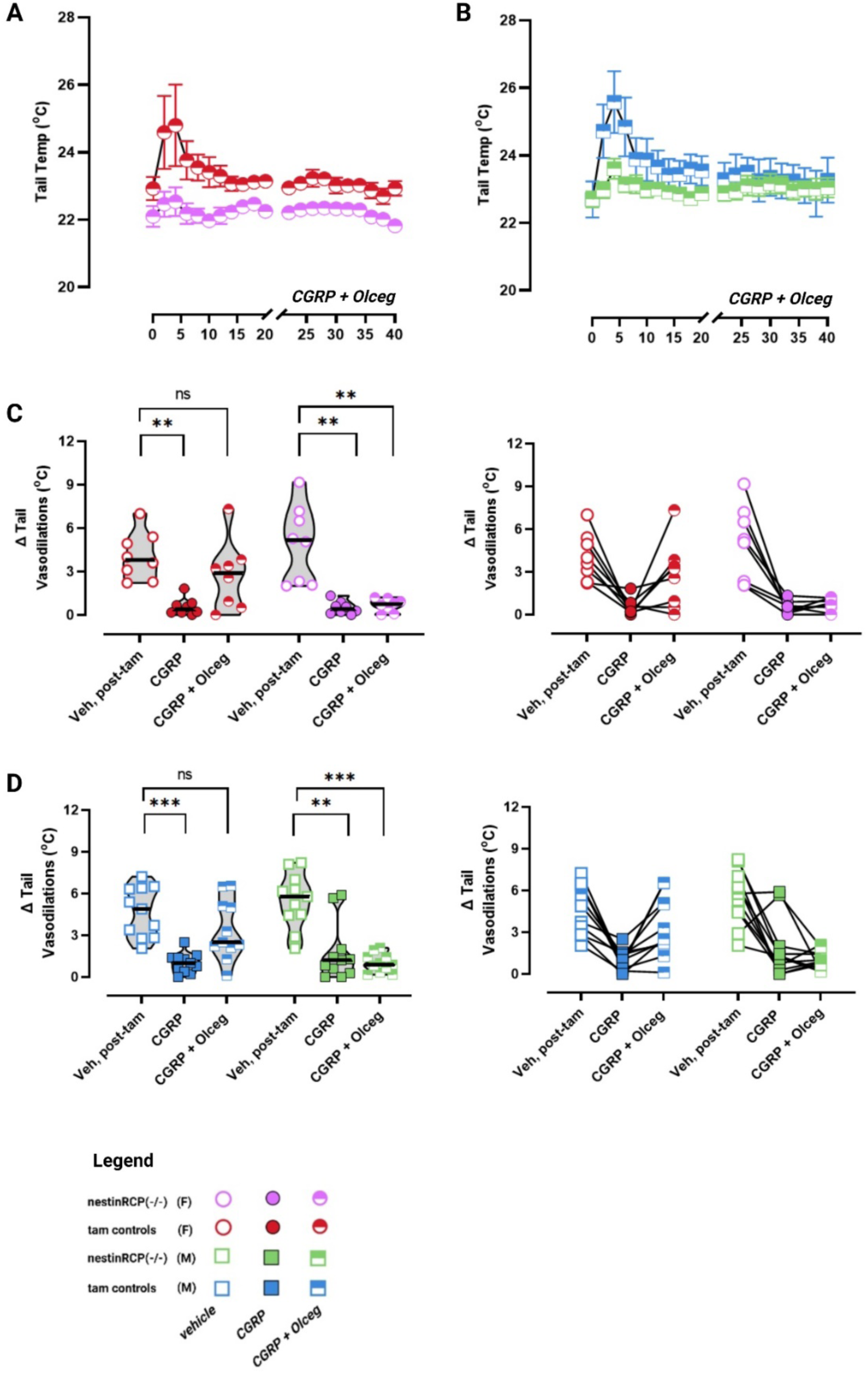
Δ tail vasodilations are compared to assess efficacy of the CGRP receptor antagonist olcegepant in blocking CGRP-induced changes. **(A-B)** Raw temperatures after CGRP + olcegepant treatment are depicted. **(C)** In female controls, CGRP caused diminished Δ tail vasodilations but co-delivery of olcegepant blocked this effect. However, female nestinRCP (-/-) did not respond to olcegepant and still exhibited diminished Δ tail vasodilations due to CGRP (D) A similar phenomenon occurred in the males, as olcegepant blocked CGRP’s effects in the male controls but male nestinRCP (-/-) was still affected by CGRP. A before/after plot is placed adjacent to violin plots to observe intra-animal changes. Full list of F-statistics and p-values can be found in Table 2. Sample sizes include 11M/8F nestinRCP (-/-) and 11M/8F controls. p < 0.05*, p < 0.01**, p < 0.001***.

In comparison, was also studied to compare with olcegepant’s effects. Raw temperatures after 1.0x CGRP + 1.0x rizatriptan are presented (**Fig. 5A-B**). In either sex, nestinRCP (-/-) and control mice did not significantly respond to rizatriptan (**Fig. 5C-D**), as Δ tail vasodilations were still diminished during 1.0x CGRP + 1.0x rizatriptan testing as observed with 1.x CGRP testing.

**Fig. 5:**
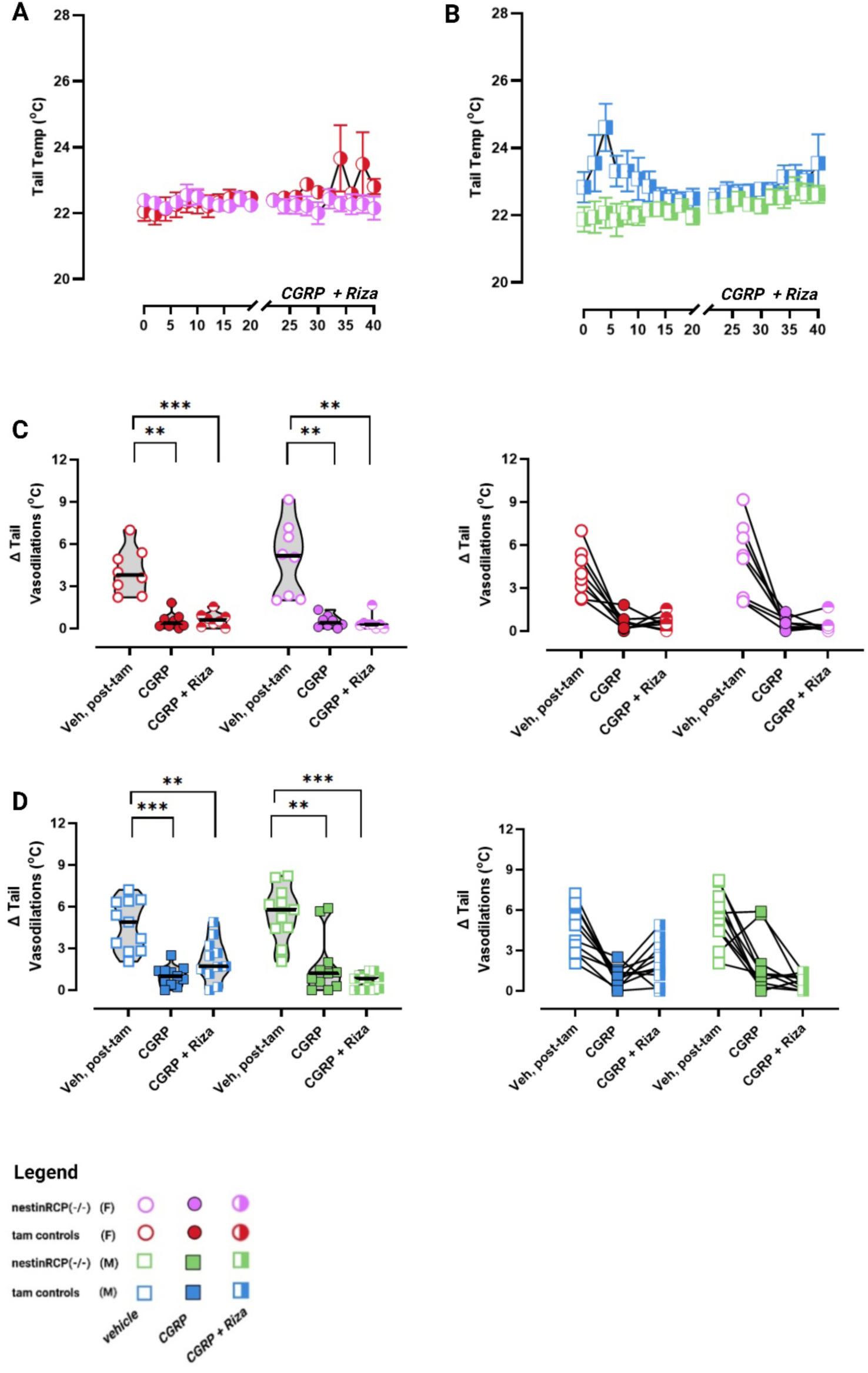
Δ tail vasodilations are compared to assess the efficacy of the serotonin receptor agonist rizatriptan in blocking CGRP-induced changes. **(A-B)** Raw temperatures after CGRP + rizatriptan treatment are depicted. **(C)** In female controls, CGRP caused diminished Δ tail vasodilations and CGRP+rizatriptan did not block these effects. **(D)** Similarly in males, rizatriptan did not block CGRP-induced changes. A before/after plot is inserted next to violin plots for intra-animal observations. F-statistics and p-values can be found in bottom half of Table 2. Sample sizes include 11M/8F nestinRCP (-/-) and 11M/8F controls. p < 0.05*, p < 0.01**, p < 0.001***

### Baseline postural sway does not differ between nestinRCP (-/-) and controls

Prior to testing CGRP and the effects of blockers, baseline CoP responses were compared between nestinRCP (-/-) and tamoxifen-treated controls. No differences were observed between controls and the nestinRCP (-/-) during vehicle-treated sway (**Fig. 6A**).

**Fig. 6:**
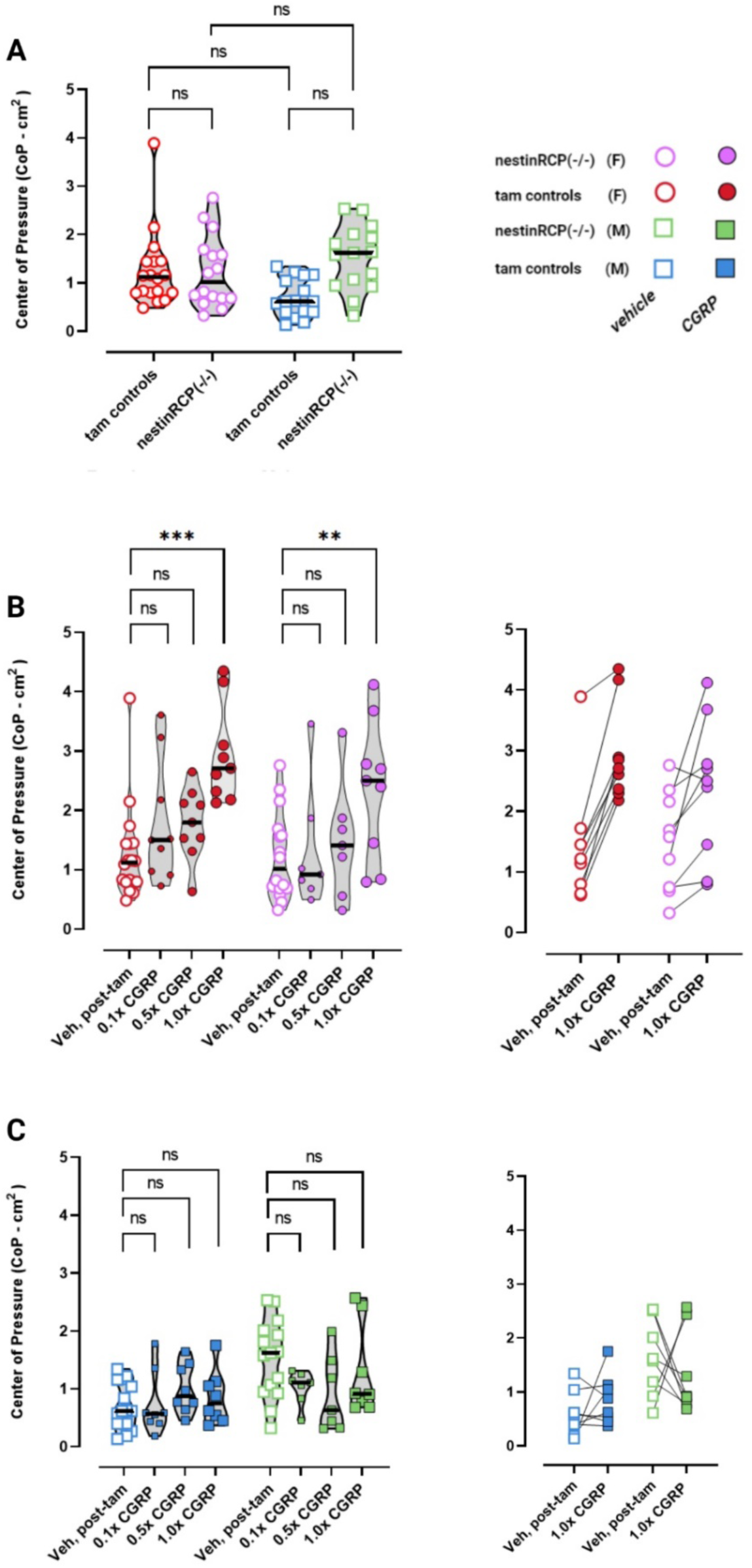
(A) 2-way ANOVA computed the factors i) male vs female and ii) controls versus nestinRCP (-/-) on sway obtained after vehicle injections (saline). Biological sex had no effect on sway (F(1, 62) = 0.75, p = 0.4), whereas a difference was observed between controls and nestinRCP (-/-) (F(1,62) = 5.05)). However, Tukey’s multiple comparisons test indicated no post- hoc comparisons to be significant. **(B)** Calcitonin gene-related peptide (CGRP) induces higher CoP ellipse areas in female tamoxifen treated controls and nestinRCP (-/-) at the 1.0x dose. **(C)** Male controls and nestinRCP (-/-) do not show changes in their CoP ellipse areas to any dose of CGRP. Before/after plots are placed adjacent to violin plots and show animals that specifically were tested with 1.0x CGRP. Sample sizes are listed: vehicle – 16M/18F nestinRCP (-/-) and 15M/16F controls; 0.1x and 0.5x CGRP -7M/7F nestinRCP (-/-) and 8M/9F controls; 1.0x CGRP - 8M/9F nestinRCP (-/-) and 8M/9F controls. Dose response statistics can be found in top half of Table 2. p < 0.05*, p < 0.01**, p < 0.001***

### Female mice exhibit significant postural sway at highest CGRP dose (0.1. mg/kg), while male mice sway is not sensitive to CGRP

Changes in center of pressure (CoP) after IP 0.1x CGRP, 0.5x CGRP, and 1.0x CGRP were examined using the same statistical tools used during motion-induced nausea studies. We observed increasing numbers of female nestinRCP (-/-) and their respective controls being sensitized to the effects of CGRP at higher doses (**Fig. 6B**). Female nestinRCP (-/-) experienced higher CoP areas at 1.0x CGRP than was measured during their vehicle test (Tukey, *adj. p = 0.009*), and this observation was similarly observed in the tamoxifen-treated controls (*Tukey, adj. p < 0.001).* In contrast, male nestinRCP (-/-) and controls did not response to increasing doses of CGRP and their CoP areas resembled their vehicle response (**Fig. 6C**).

### Olcegepant effectively blocks CGRP-induced sway increases in a female control majority, but lacks efficacy in a subset of nestinRCP (-/-)

Due to the greater CoP areas observed at 1.0x CGRP in female mice, we further examined if migraine blockers olcegepant and rizatriptan can ameliorate changes in sway induced by 1.0x CGRP. We observed that CGRP+olcegepant reduced sway in all female controls, but a subset of female nestinRCP (-/-) did not respond to olcegepant, as clearly seen in the before-after plots (**Fig. 7A**). Postural sway measured in female nestinRCP (-/-) during CGRP+olcegepant testing resembled CGRP only testing. Similar findings were observed during testing after IP delivery of CGRP + rizatriptan. Female controls exhibited reduced CoP area when treated with CGRP + rizatriptan compared to their CGRP only test, whereas female nestinRCP (-/-) did not appear to respond to rizatriptan (**Fig. 8A**). In contrast, male nestinRCP (-/-) and controls did not respond to olcegepant (**Fig. 7B**) or rizatriptan (**Fig. 8B)** during CoP testing.

**Fig. 7:**
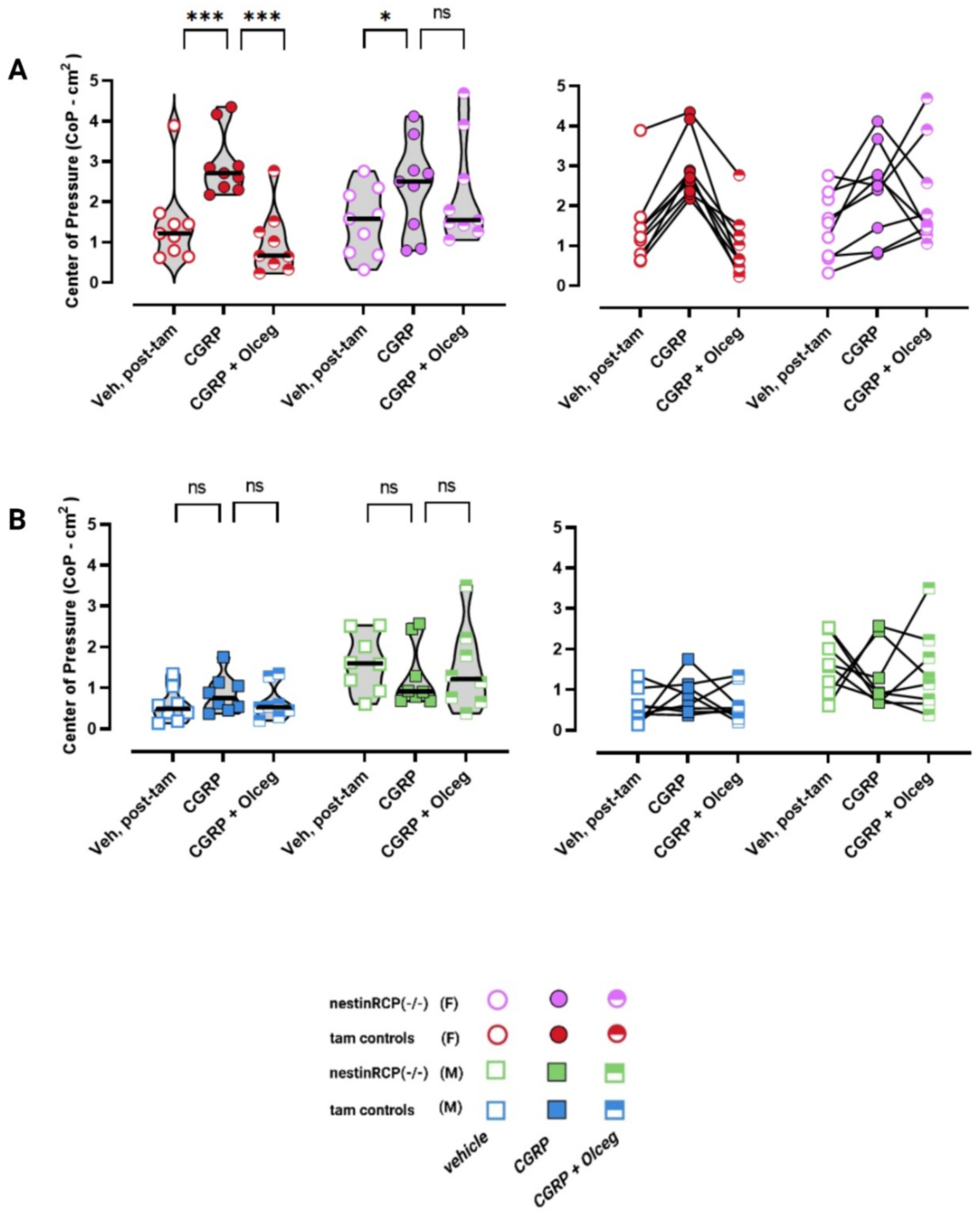
Changes in CoP ellipse areas are observed after vehicle, CGRP, and CGRP + olcegepant treatment in controls and nestinRCP (-/-). **(A)** In female controls, CGRP causes increases in postural sway but co-administration of olcegepant blocks CGRP-induced changes. In contrast, a subset of female nestinRCP (-/-) do not respond to olcegepant and still exhibit postural sway that resembles CGRP testing. **(B)** Male mice do not significant change their sway in response to CGRP or CGRP + olcegepant. Before/after plots are adjacent to violin plots. Sample sizes include 8M/9F nestinRCP (-/-) and 8M/9F controls. Full list of F and p-values are found in the middle portion of Table 2. p < 0.05*, p < 0.01**, p < 0.001***, p < 0.0001****

**Fig. 8:**
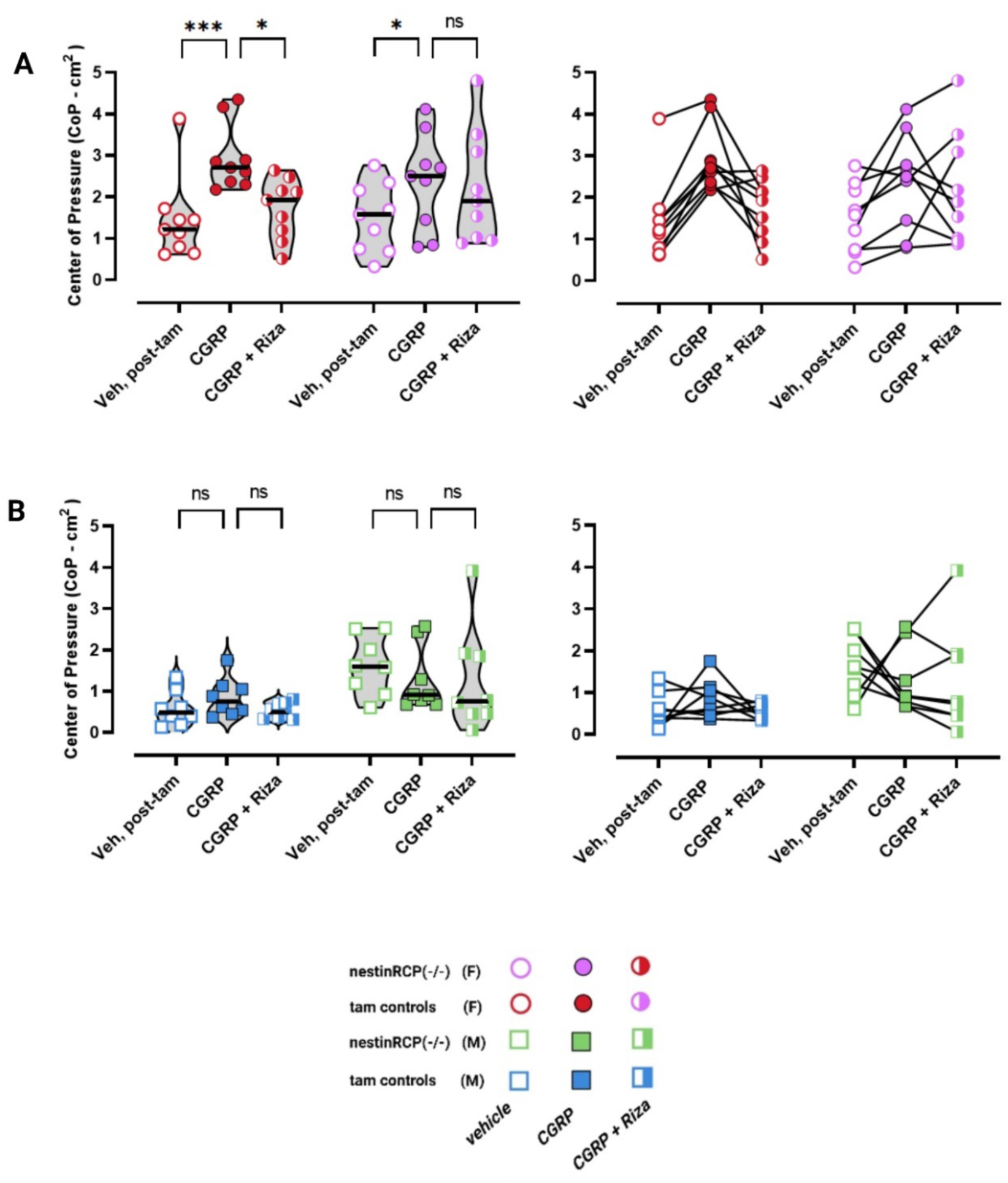
Changes in CoP ellipse areas are observed after vehicle, CGRP, or CGRP + rizatriptan treatment in controls and nestinRCP (-/-). **(A)** Similar to olcegepant studies, rizatriptan was shown to block CGRP-induced changes in sway for female controls, but not in the female nestinRCP (-/-). **(B)** Male mice do not respond to CGRP or CGRP+rizatriptan. Sample sizes include 8M/9F nestinRCP (-/-) and 8M/9F controls. Full list of F and p-values are found in the middle portion of Table 2. p < 0.05*, p < 0.01**, p < 0.001***, p < 0.0001****

**Table 2** is provided with details regarding multivariate statistics used in both assays, and their respective F-statistics and p-values.

## Discussion

Supplementing prior research where we studied the effects of systemic CGRP in motion-induced nausea and postural sway in C57BL6/J mice, this study assessed CGRP’s effects on these behaviors in a mouse model with RCP loss in primarily the nervous system. We were presented with several key findings.

First, systemic CGRP still leads to changes in postural sway and motion-induced nausea in nestin RCP (-/-) that resembles littermate controls – suggesting neural RCP loss does not impact the occurrence of these behaviors. Female nestinRCP (-/-) and their tamoxifen-treated controls still present with higher sway after administration of 0.1 mg/kg CGRP, whereas male nestinRCP (-/-) and controls do not modulate their sway to systemic CGRP. Systemic CGRP also diminishes Δ tail vasodilations in the nestinRCP (-/-) and tamoxifen-treated controls, which was also observed in C57BL6/J not treated with tamoxifen [19, 20]. It is generally understood that RCP is not required for CGRP to bind to the CLR/RAMP1 receptor, but these findings suggest that downstream signaling that is relevant for these behaviors still continues even with RCP loss.

While CGRP induced changes in the nestinRCP (-/-) that resembled tamoxifen-treated controls and in our prior study with wildtype C57BL6/KJ, the scope of CGRP’s impact on nausea responses in these rodent models is still unclear. The selective knockout of RCP in nestin-cells suggests changes in the signaling of not just the nervous system, but potentially endothelium and smooth muscle[21]. Prior literature has shown that CGRP induces smooth muscle relaxation and that RCP is implicated in smooth muscle contractility, so RCP loss in smooth muscle may impact gastrointestinal physiology and peristalsis activity, which may signal to the rodent area postrema and impact nausea response. A future direction requires assessing other nausea correlates in mice, such as behavioral taste aversion. Interestingly, the activation of

Gαs-coupled signaling in GLP1R neurons located in the area postrema is sufficient to condition taste aversion, and the area postrema houses CGRP receptors [22, 23]. Taste aversion for nausea studies in the nestinRCP (-/-) is a future experiment for consideration.

In cell culture studies, RCP has been demonstrated to promote the coupling of the CGRP receptor and a number of B family GPCRs to cAMP using Gαs. Interestingly, RCP is not associated with pERK or Gq/Ca^2+^ activation [13], and so CGRP-induced behaviors in the nestinRCP (-/-) mouse may be attributed to alternate downstream pathways that do not require RCP – such as mitogen activated protein kinase (MAPK), Gαq or Gα independent signaling. Gα independent signaling has been attributed to nitric oxide release, and we have previously shown that intraperitoneally (IP) delivered sodium nitroprusside – a nitric oxide donor - leads to diminished Δ tail vasodilations in the C57BL/6J much like is observed after IP CGRP [20]. Thus, these alternate pathways may explain CGRP-induced behaviors despite RCP loss.

The main findings of this study allude to a potential change in the CGRP receptor conformation due to RCP loss, that manifests into behavioral changes when its selective knocked out in the nervous system. In the motion-induced thermoregulation studies, Δ tail vasodilations were recovered in tamoxifen-treated males and less frequently in females with the co-administration of olcegepant. This observation is internally consistent as we saw similar results in prior motion-induced thermoregulation studies of the C57BL/6J. However, the nestinRCP (-/-) mice did not respond to olcegepant. This observation carried over to the center of pressure testing, where olcegepant was able to reduce CoP areas in tamoxifen-treated females and counter CGRP-induced changes, but olcegepant was less effective in the female nestinRCP (-/-). A unique finding in this study’s sway experiments is the presentation of a blocker effect by rizatriptan, which was not seen in the motion-induced nausea tests. Triptans like rizatriptan are known to be poor drugs for vestibular migraine’s vertigo, but still exert an antimigraine effect by potentially reducing circulating CGRP via a number of proposed mechanisms [24, 25] Olcegepant was previously thought as a highly specific and potent nonpeptide antagonist to the CGRP and adrenomedullin receptors [26], and its affinity for CGRP receptors is species dependent – with a higher affinity for the human receptor over the mouse receptor. Prior research has suggested olcegepant blocks CGRP signaling by attenuating cAMP accumulation and phosphokinase A (PKA)-dependent pain responses in neurons. Our results suggest RCP loss would lessen olcegepant’s efficacy as an antagonist, if its activity was dependent on RCP coupling of cAMP to the receptor. We propose that in the presence of RCP, olcegepant binds to the receptor and antagonizes CGRP signaling mediated by cAMP. However, in the absence of RCP, olcegepant may lose its affinity for the receptor. Thus, while the activity of the CGRP peptide requires CLR/RAMP1, the efficacy of antagonists like olcegepant may require CLR/RAMP1/RCP, thereby explaining the results of this study. An interesting future direction for addressing RCP’s role in olcegepant’s action is to test MAPK and other Gα independent activity *in vitro*, using a tamoxifen-inducible cell culture line.

Molecular docking and molecular dynamics simulations have attempted to model the drug-receptor complex and analyze interactions between the CGRP receptor and the gepants, and have discovered subtle differences in the binding of ubrogepant and rimegepant from olcegepant [27]. A future direction would be to further characterize the gepants in our motion-induced nausea and postural sway assays, and observe if other gepants resembles current data in olcegepant, as these subtle differences may be linked to RCP’s presence. Despite these claim, it is unclear how RCP factors into molecular docking and dynamics simulations, because prior studies have emphasized CGRP interactions with CLR/RAMP1 [11].

The current results are thought provoking, but it is still unclear how RCP loss factors into desensitization and internalization of the CGRP receptor, as a potential change in the receptor conformation due to RCP loss may influence long-term response to CGRP and similar ligands. In addition, while the focus of the study has been on primarily CGRP-RCP interactions, it is also necessary to recognize that RCP is involved in other GPCRs and that CGRP may exhibit cross- talk at amylin (AMY) and adrenomedullin (ADM) receptors. While RCP is a component of the

ADM receptor, amylin is free from it [11, 28]. It is of interest to gauge levels of AMY signaling after RCP loss in different cell types, as one could hypothesize that reduced CGRP/ADM signaling may leads to a compensatory change in the cell via upregulated AMY signaling. While not a focus of this paper, this hypothesis should be explored *in vitro*.

This work is part of a greater picture to understand how RCP and related cytosolic, peripheral membrane proteins may be modulated for selective activation of GPCR signaling, in an effort to elucidate disease mechanisms and devise alternate therapeutic strategies for migraine treatment and related indications.

## Conflict of interest statement

The authors declare no competing financial interests.

### Acknowledgments

This work was fully supported by NIH R01DC017261 (AEL). We would like to thank Abigail Dweh and Anna Guo for their contributions to the data analysis.

---

**Supplementary Fig. S1:**
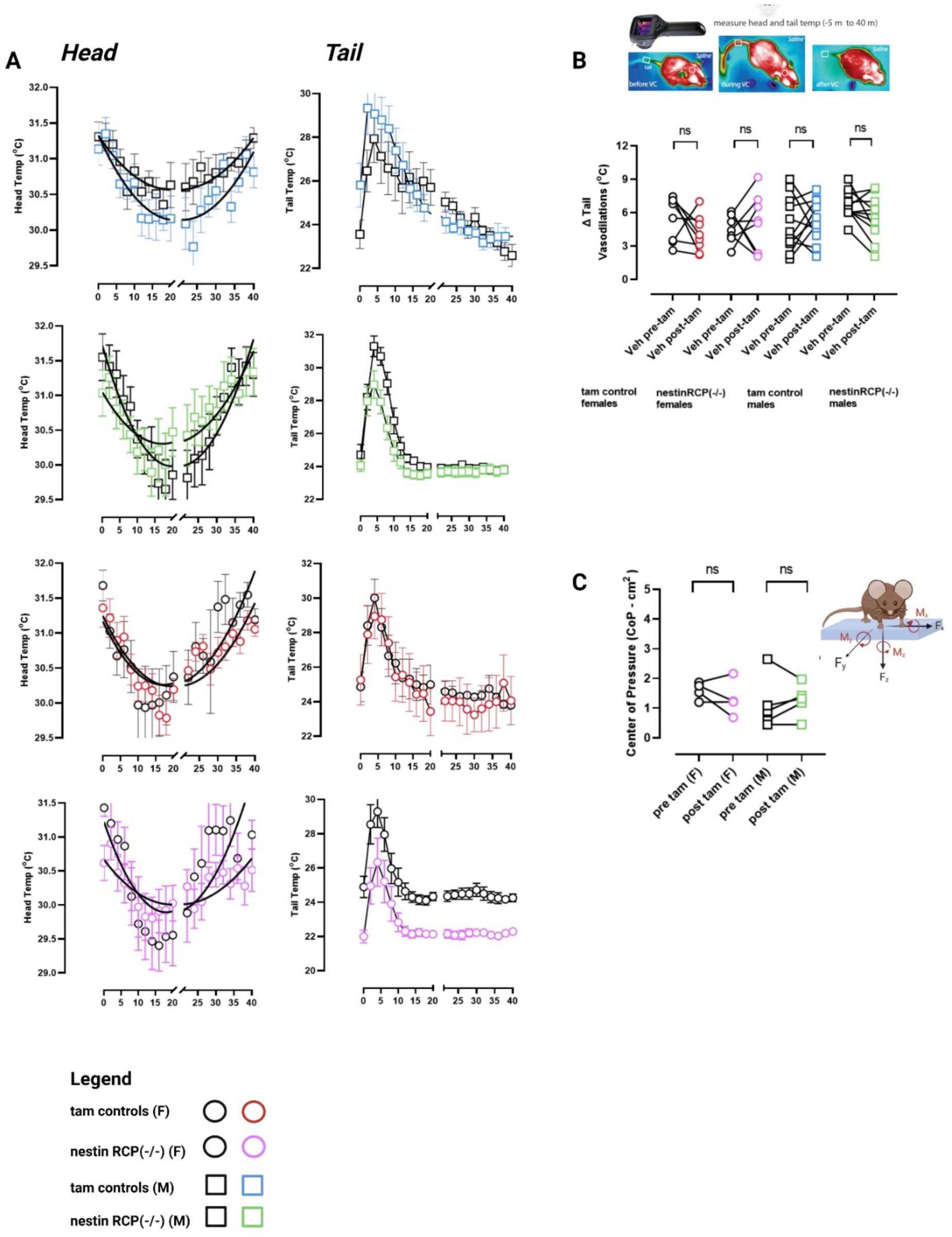
(A) Head and tail temperatures are shown in tamoxifen-treated controls and nestinRCP (-/-) before and after tamoxifen induction protocol. (B) Tamoxifen treatment does not impact the presentation of Δ tail vasodilations (n = 11M/8F nestinRCP (-/-) and 11M/8F controls). (C) Tamoxifen treatment does not impact postural sway in the nestinRCP (-/-), as baseline center of pressure ellipse areas are similar in mice pre- and post-tamoxifen (n = 4F/ 5M).

